# BNCT Triggers Immunogenic Cell Death in B16 Melanoma Without Accompanying T-Cell Infiltration—Implications for Combination Immunotherapy

**DOI:** 10.64898/2026.11.04.764456

**Authors:** Rui Quan, Hefa Huang, Lili Zhang, Chengyu Zhang, Shuyu Chang, Menglin Wu, Shixu Zhang, Xingkang Su, Lu Zhang, Zhongying Dai, Long Gu

## Abstract

**Objective:** Boron neutron capture therapy (BNCT) is a targeted radiotherapy with highly selective tumor killing; its potential to induce immunogenic cell death (ICD) and modulate the tumor immune microenvironment has attracted attention. However, whether BNCT-induced immune activation can overcome the T-cell exclusion barrier of immunologically “cold” tumors and translate into intratumoral effector T-cell infiltration remains unclear.

**Methods:** A B16 melanoma model was established in C57BL/6 mice. Mice were assigned to CK, N10/N20 (neutron irradiation alone for 10 or 20 min), BPA, and BNCTN10/BNCTN20 groups (BPA 500 mg/kg plus 10 or 20 min irradiation) to evaluate tumor suppression and survival. In a separate cohort (CK, N20, and BNCTN20), the most efficacious regimen (BNCTN20) was compared with CK on days 3 and 7: H&E staining; immunohistochemistry/immunofluorescence for γH2AX, HMGB1, cleaved caspase-3, PD-L1, CD8⁺, and granzyme B; ELISA for cytokines and chemokines; and ICP-OES for ¹⁰B distribution.

**Results:** BNCT significantly suppressed tumor growth. When CK tumors reached the humane endpoint (≈2000 mm³), N20 tumor volume was half that of CK (TGI ≈ 49.8%), whereas BNCTN20 reached only one tenth (TGI ≈ 88.9%, p < 0.001), with prolonged survival (p < 0.001). BNCT induced γH2AX-associated DNA damage, HMGB1 release, and cleaved caspase-3-mediated apoptosis; upregulated PD-L1; induced a Th1-type response (elevated IFN-γ/TNF-α, reduced IL-10); and enhanced CXCL9/10 signaling. However, intratumoral CD8⁺ T-cell infiltration and granzyme B expression did not increase, indicating that this activation was not translated into a local effector T-cell response.

**Conclusions:** BNCT inhibited tumor growth and prolonged survival while inducing ICD and a systemic Th1 response; nevertheless, systemic activation failed to translate into local CD8⁺ T-cell infiltration, while PD-L1 was upregulated—a “strong direct killing but blocked immune translation” profile in the B16 cold tumor. PD-L1-associated adaptive immune resistance may limit this conversion, providing rationale for combining BNCT with immune checkpoint blockade.

## Introduction

Malignant melanoma (MM) is one of the most aggressive cutaneous malignancies, with a high metastatic propensity and poor prognosis. In recent years, the clinical application of immune checkpoint inhibitors (ICIs) has markedly improved outcomes in advanced melanoma, raising the 5-year overall survival rate from less than 20% to approximately 35%–40%^1–3^. However, a substantial proportion of patients still fail to derive durable benefit from ICIs, particularly those with immunologically “cold tumors” that respond poorly to programmed death-1/programmed death-ligand 1 (PD-1/PD-L1) pathway blockade^1^. B16 melanoma is a classic immunologically cold tumor model characterized by low tumor immunogenicity, scarce tumor-infiltrating lymphocytes, and refractoriness to single-agent immune checkpoint blockade, making it a canonical model for evaluating whether novel immunotherapeutic strategies can turn “cold” tumors “hot”^4,5^.

BNCT is a binary targeted radiotherapy modality. The ¹⁰B-enriched compound L-4-boronophenylalanine (BPA) is selectively accumulated in tumor tissue via L-type amino acid transporter 1 (LAT1), which is highly expressed on tumor cells; subsequent epithermal neutron irradiation triggers the ¹⁰B(n,α)⁷Li nuclear reaction, generating high linear energy transfer (high-LET) α particles and ⁷Li heavy ions. Because the tissue range of both particles is only approximately 5–9 μm, comparable to the diameter of a single cell—BNCT selectively kills boron-loaded tumor cells while minimizing damage to surrounding normal tissue^6,7^. In recent years, BNCT has shown unique advantages in the treatment of various refractory tumors. In melanoma in particular, preclinical experiments and early clinical explorations have confirmed its strong potential for tumor control, offering a new strategy to overcome resistance to conventional radiotherapy^8^.

The immunomodulatory effects of BNCT are receiving increasing attention. Studies have shown that, compared with an equivalent dose of X-rays, BNCT causes less damage to immune cells and induces immunogenic cell death (ICD), thereby improving the tumor immune microenvironment by promoting the activation of dendritic cells (DCs), T cells, and NK cells^9,10^. However, ICD induced by BNCT alone is difficult to translate into sustained, systemic antitumor immunity, and abscopal effects have rarely been reported; combination with immune adjuvants or immune checkpoint blockade is expected to further enhance its therapeutic efficacy.^11^ In melanoma models, BNCT combined with anti-PD-L1 therapy enhanced intratumoral T-cell infiltration and activation; in addition, boron neutron immunotherapy (B-NIT) showed the potential to induce abscopal effects and improve treatment responses in immunotherapy-resistant melanoma models^5,12^. Nevertheless, whether single-agent BNCT can breach the T-cell exclusion barrier in the classic B16 “cold tumor” model, whether the immune activation it induces can be effectively converted into intratumoral effector T-cell infiltration, and the role of PD-L1 in this process remain unsupported by systematic evidence. Using a C57BL/6 mouse B16 melanoma model, this study first evaluated the antitumor efficacy and survival benefit of BNCT at different neutron doses, and then compared the optimal treatment group (BNCTN20) with the control group (CK) in mechanistic analyses of ICD-related molecules, immune cytokines, chemokines, T-cell infiltration, and PD-L1 expression, combined with assessment of BPA tissue distribution in a bilateral tumor model, to delineate the immune activation profile of BNCT in cold tumors and its potential limiting mechanisms.

## Materials and Methods

### Cell Culture

B16 murine melanoma cells (BeNa Culture Collection, BNCC) were cultured in DMEM supplemented with 10% fetal bovine serum and 1% penicillin–streptomycin at 37 °C in a humidified 5% CO₂ atmosphere.

### Animals and Tumor Models

Female C57BL/6 mice (6–8 weeks old, 20 ± 2 g; Guangdong Yaokang Biotechnology Co., Ltd.) were housed under specific pathogen-free (SPF) conditions. All animal experiments were approved by the Ethics Committee of the Putian Lanhai Nuclear Medicine Research Center (approval No. 2026(01)) and conducted in accordance with relevant animal experiment regulations.

For the unilateral tumor model, B16 cells (5 × 10⁵ cells in 100 μL PBS) were inoculated subcutaneously into the right dorsal flank. For the bilateral tumor model, B16 cells (5 × 10⁵ and 3 × 10⁵ cells) were inoculated into the right dorsal flank and left shoulder, respectively. Tumor volume was calculated as *V* = 0.5 × length × width².

### Neutron Irradiation and BPA Administration

BNCT experiments were performed using the accelerator-based BNCT system at Lanzhou University (Fig. 1a–c). The proton beam energy was 2.6 MeV with a beam current of 6 mA, and the epithermal neutron flux at 2 cm from the beam exit was 6.96 × 10⁸ n·cm⁻ ²·s⁻ ¹ (calibrated by the activation method). BPA (L-¹⁰B-BPA lyophilized powder, provided by Shenzhen Zhonghe Headway Biotechnology Co., Ltd.) was dissolved in sterile water to prepare a 30 mg/mL working solution. Mice received tail-vein injection of BPA (500 mg/kg) and were irradiated 1 h later.

**FIG. 1.**
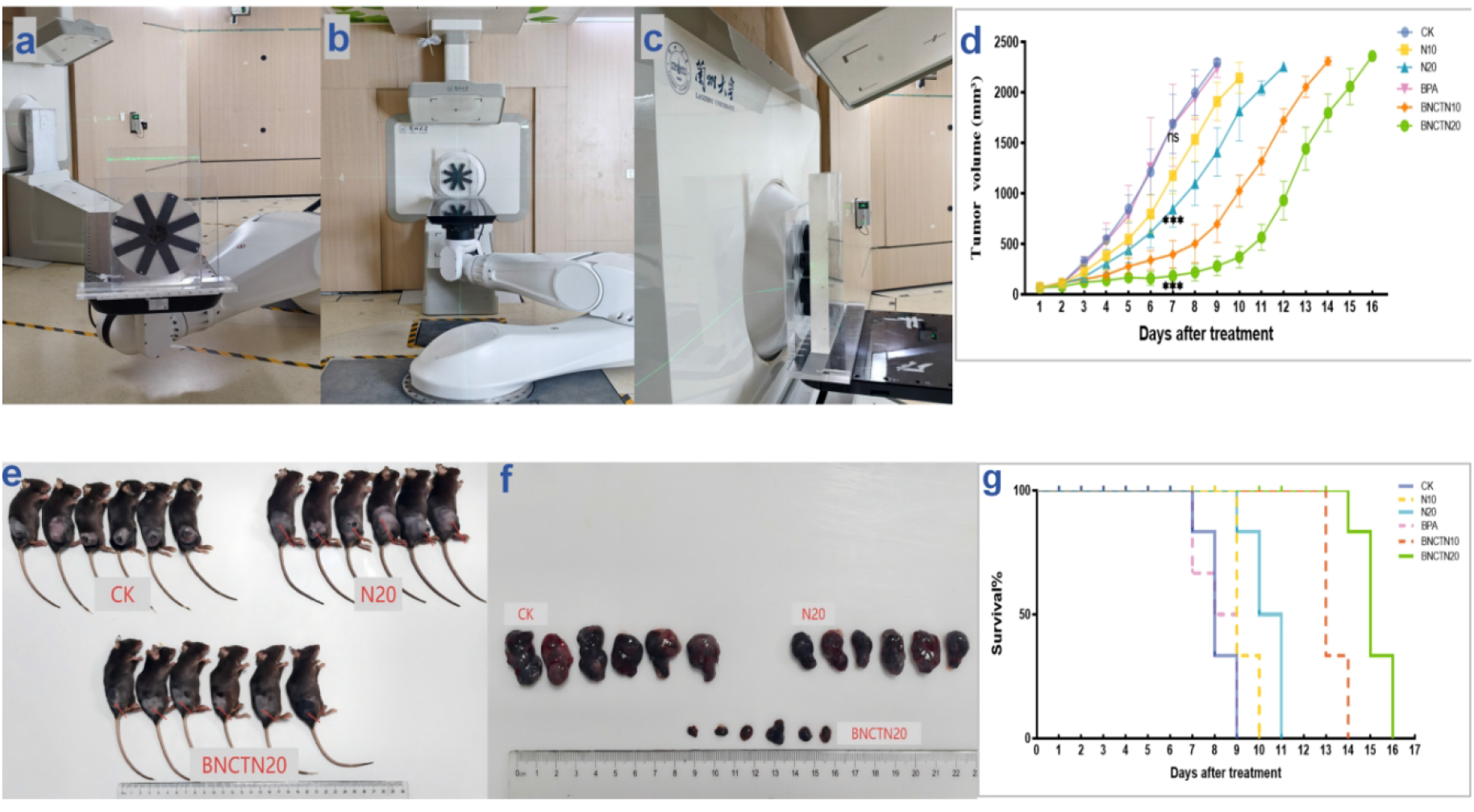
BNCT irradiation setup and tumor growth inhibition in B16 melanoma-bearing mice. (a) Mouse immobilization and positioning device for BNCT; (b) overall layout of the irradiation apparatus (beam collimator, robotic arm, and irradiation platform); (c) beam exit and positioning laser, with the tumor 2 cm from the beam port. (d) Tumor growth curves of each group (n = 6). Intergroup comparisons on day 7 were performed using one-way ANOVA with Tukey’s multiple comparisons test: the BPA group did not differ significantly from CK (ns); the N20 and BNCTN20 groups differed significantly from CK, and BNCTN20 differed significantly from N20 (all *p* < 0.001). Data are presented as mean ± SD. (e) Photographs of body-surface tumors on day 7 (from the independent tissue-collection cohort). (f) Photographs of excised tumors (from the same cohort). (g) Kaplan–Meier survival curves; comparisons among groups were performed using the log-rank (Mantel–Cox) test.

### Experimental Grouping and Design

When tumor volumes reached 60–80 mm³, mice were randomly assigned to groups.

### Efficacy Evaluation Experiment

Mice were randomly divided into CK, N10, N20, BPA, BNCTN10, and BNCTN20 groups (n = 6 per group). The CK group received no treatment; the BPA group received tail-vein injection of BPA (500 mg/kg) without irradiation; the N10/N20 groups received neutron irradiation alone for 10 or 20 min, respectively; and the BNCTN10/BNCTN20 groups received neutron irradiation for 10 or 20 min, respectively, 1 h after tail-vein injection of BPA (500 mg/kg). During the experiment, animal status was monitored and tumor volumes were measured daily, tumor growth curves were plotted, and survival was recorded. Humane endpoints were defined as tumor volume ≥ 2000 mm³, tumor ulceration, or signs such as cachexia, restricted mobility, or loss of appetite indicating inability to maintain normal vital activities, at which point euthanasia was performed. Tumor growth inhibition (TGI) was calculated as TGI (%) = (1 – V_treatment_ / V_CK_) × 100%.

### Histological and Mechanistic Study Experiment

In a separate cohort, mice were randomly divided into CK, N20, and BNCTN20 groups (n = 12 per group). On days 3 and 7 after irradiation, 6 mice per group were randomly selected for retro-orbital blood collection to prepare serum, followed by euthanasia. Body-surface and excised tumor images were photographed. Tumor tissues from the CK and BNCTN20 groups were used for mechanistic analyses, while the N20 group served as a neutron-irradiation control for morphological comparison. Tumor tissues were fixed in 4% paraformaldehyde and embedded in paraffin for H&E staining, immunohistochemistry, and immunofluorescence.

### H&E Staining

Paraffin sections were stained with hematoxylin and eosin (H&E), and tumor necrosis, apoptosis, and morphological changes were observed under a light microscope.

### Immunohistochemistry (IHC)

After antigen retrieval and blocking, paraffin sections were incubated overnight at 4 °C with the following primary antibodies: anti-CD8α (Abcam, 1:100), anti-granzyme B (Proteintech, 1:50), anti-PD-L1 (Proteintech, 1:100), and anti-cleaved caspase-3 (Proteintech, 1:100), followed by incubation with HRP-conjugated secondary antibodies, DAB chromogenic development, and hematoxylin counterstaining. Integrated optical density (IOD) and positive area were measured using ImageJ, and IOD/Area was calculated.

### Immunofluorescence (IF)

After antigen retrieval and blocking, paraffin sections were incubated overnight at 4 °C with the following primary antibodies: anti-γH2AX (Proteintech; 1:100) and anti-HMGB1 (Proteintech; 1:100), followed by incubation with CoraLite 594-conjugated goat anti-rabbit IgG (H+L) secondary antibody (Proteintech; 1:100) for 1 h at room temperature in the dark, and DAPI counterstaining of nuclei. Images were acquired by fluorescence microscopy and quantified using ImageJ.

### Serum Cytokine ELISA

After retro-orbital blood collection, samples were allowed to stand at room temperature for 30 min and centrifuged at 3000 rpm for 15 min at 4 °C to collect serum. Serum levels of IFN-γ (E-EL-M0048), TNF-α (E-EL-M3063), IL-10 (E-HSEL-M0004), CXCL9 (E-EL-M3077), and CXCL10 (E-EL-M0021) were measured using commercial ELISA kits (Elabscience, Wuhan, China) according to the manufacturer’s instructions.

### Boron Biodistribution Analysis by ICP-OES

For boron biodistribution analysis, bilateral tumor-bearing mice received tail-vein injection of BPA (500 mg/kg), and blood, bilateral tumors, and major normal tissues (brain, heart, liver, lung, spleen, kidney, skin, and muscle) were collected at 0.5, 1, 1.5, 2, 2.5, 3, 4, 5, 6 and 7 h after administration (n = 4 per time point). Tissue samples were weighed and digested in concentrated nitric acid at 70 °C for 6 h. The digested solutions were diluted to a fixed volume with ultrapure water, and ^10^B concentrations were determined by inductively coupled plasma optical emission spectrometry (ICP-OES; PQ 9100 Elite, Analytik Jena, Germany). Tumor-to-blood (T/B) and tumor-to-normal tissue (T/N) ratios were calculated from the measured concentrations.

### Statistical Analysis

Data are presented as mean ± SD. Comparisons between two independent groups were performed using Welch’s t-test; comparisons between bilateral tumors and paired tissues from the same mouse were performed using paired t-tests; comparisons among multiple groups were performed using one-way ANOVA with Tukey’s post-hoc test; tumor growth curves were analyzed by two-way repeated-measures ANOVA with correction for multiple comparisons; survival was analyzed by the Kaplan–Meier method with the log-rank test. A *p* value < 0.05 was considered statistically significant.

## Results

### BNCT Inhibits B16 Melanoma Growth and Prolongs Survival of Tumor-Bearing Mice

#### Cohort 1: Efficacy Evaluation

In the efficacy evaluation experiment, when the first mouse in the CK group reached the humane endpoint (day 7, ≈2000 mm³), tumor growth was analyzed across groups. Tumor volume in the N20 group was 50.2% of that in the CK group (TGI ≈ 49.8%), whereas the BNCTN20 group reached only 11.1% of CK (TGI ≈ 88.9%) (Fig. 1d); the BPA group did not differ significantly from CK, while all other intergroup comparisons were statistically significant (all *p* < 0.001). Kaplan–Meier survival analysis further showed (Fig. 1g) that median survival was 8 days in the CK group and 10.5 days in the N20 group, and was further prolonged to 15 days in the BNCTN20 group, significantly longer than both the CK and N20 groups (both *p* < 0.001).

#### Cohort 2: Tumor Morphology Observation

In the independent tissue-collection cohort, mice were euthanized on days 3 and 7 after irradiation (n = 6 per group per time point). On day 7, body-surface and excised tumors were photographed and tumor tissues were collected. Compared with the CK and N20 groups, both the body-surface tumor burden and the excised tumor size were markedly reduced in the BNCTN20 group (Fig. 1e, f), consistent with the tumor growth curves.

### BNCT Induces Histological Damage and Apoptosis in B16 Melanoma

H&E staining on day 7 showed that tumor cells in the CK group were densely arranged with intact architecture and no obvious necrosis (Fig. 2a), whereas tumors in the BNCTN20 group exhibited marked structural disruption, extensive necrotic areas, abundant apoptotic cells, stromal congestion, and a small amount of inflammatory cell infiltration (Fig. 2b). Cleaved caspase-3 immunohistochemistry on day 7 showed markedly stronger positive staining in the BNCTN20 group than in the CK group (Fig. 2c–d). Quantitative analysis confirmed that cleaved caspase-3 expression in the BNCTN20 group was significantly increased to 2.64-fold that of the CK group (*p* < 0.01) (Fig. 2e).

**FIG. 2.**
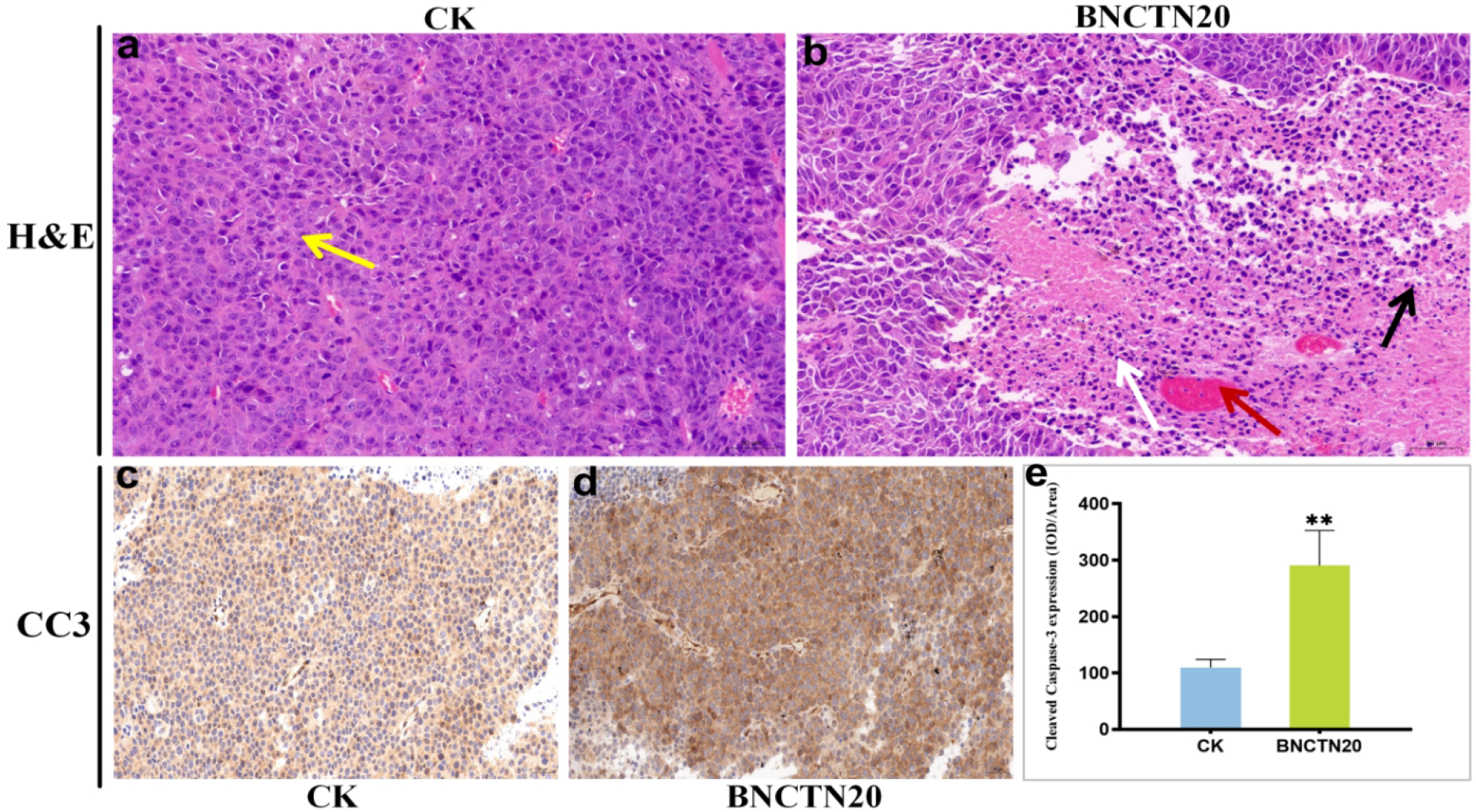
BNCT induces histological damage and apoptosis in B16 melanoma. H&E staining of tumor tissues on day 7 after irradiation: (a) CK group; yellow arrow: densely arranged tumor cells. (b) BNCTN20 group; white arrows: apoptotic cells; red arrow: stromal congestion; black arrow: inflammatory cell infiltration. (c–e) Cleaved caspase-3 immunohistochemical staining and quantitative analysis (brown-yellow indicates positive signal; blue indicates hematoxylin-counterstained nuclei; expressed as IOD/Area).

Comparisons were performed using Welch’s t-test.

### BNCT Induces DNA Damage and ICD-Related Changes

Immunofluorescence showed that on days 3 and 7 after irradiation, γH2AX red fluorescence signals in tumors of the BNCTN20 group were markedly stronger than those in the CK group at the same time points, and fluorescence intensity increased further over time (Fig. 3a). Quantitative analysis showed (Fig. 3c) that γH2AX expression in the BNCTN20 group was significantly higher than in the CK group on day 3 (*p* < 0.05), and the difference was even more pronounced on day 7 (*p* < 0.001). Within the BNCTN20 group, expression on day 7 was significantly higher than on day 3 (*p* < 0.01), indicating time-dependent accumulation, whereas no significant difference was observed between days 3 and 7 in the CK group (*p* > 0.05).

**FIG. 3.**
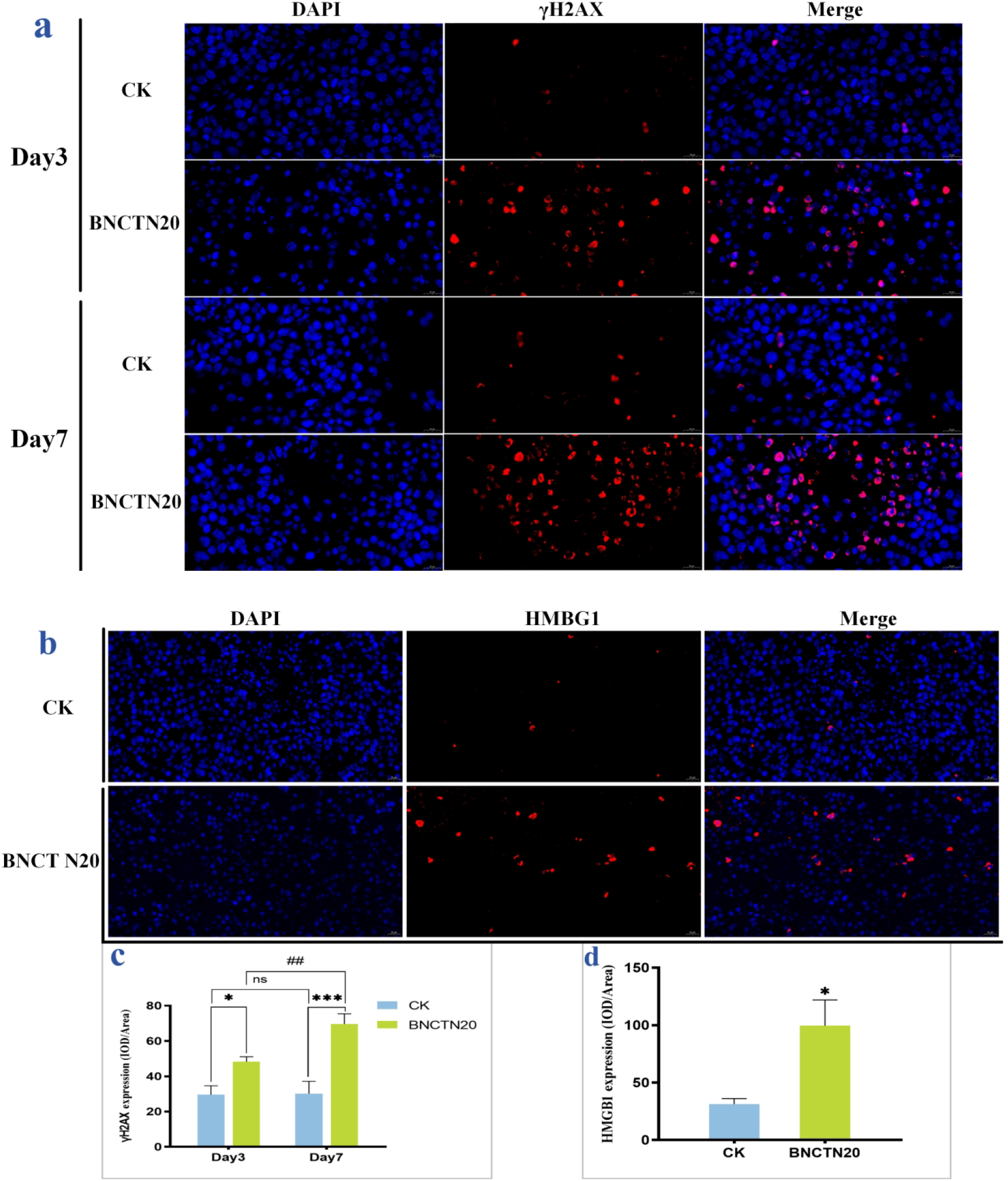
BNCT increases γH2AX expression and promotes HMGB1 release. (IF: red, γH2AX or HMGB1; blue, DAPI; Merge, overlaid images. Two-group comparisons were performed using Welch’s t-test; multi-group comparisons were performed using one-way ANOVA with Tukey’s multiple comparisons test.) (a) γH2AX immunofluorescence images of the CK and BNCTN20 groups on days 3 and 7 after irradiation. (b) HMGB1 immunofluorescence images on day 3. (c) Quantitative analysis of γH2AX fluorescence intensity (IOD/Area). (d) Quantitative analysis of HMGB1 fluorescence intensity (IOD/Area).

HMGB1 immunofluorescence staining and quantitative analysis on day 3 showed that HMGB1 red fluorescence in tumors of the BNCTN20 group was significantly stronger than in the CK group (3.18-fold, *p* < 0.05), with part of the signal located outside the nucleus, suggesting HMGB1 translocation and release from the nucleus into the cytoplasm/extracellular space (Fig. 3b, d).

### BNCT Induces a Systemic Th1 Immune Response and Enhances Chemotactic Signaling

ELISA on day 7 showed that, compared with the CK group, serum IFN-γ and TNF-α levels in the BNCTN20 group were significantly increased to 3.71-fold and 2.00-fold those of CK, respectively (both *p* < 0.05; Fig. 4a, b), whereas the level of the immunosuppressive cytokine IL-10 decreased to 0.47-fold that of CK (*p* < 0.05; Fig. 4c).

**FIG. 4.**
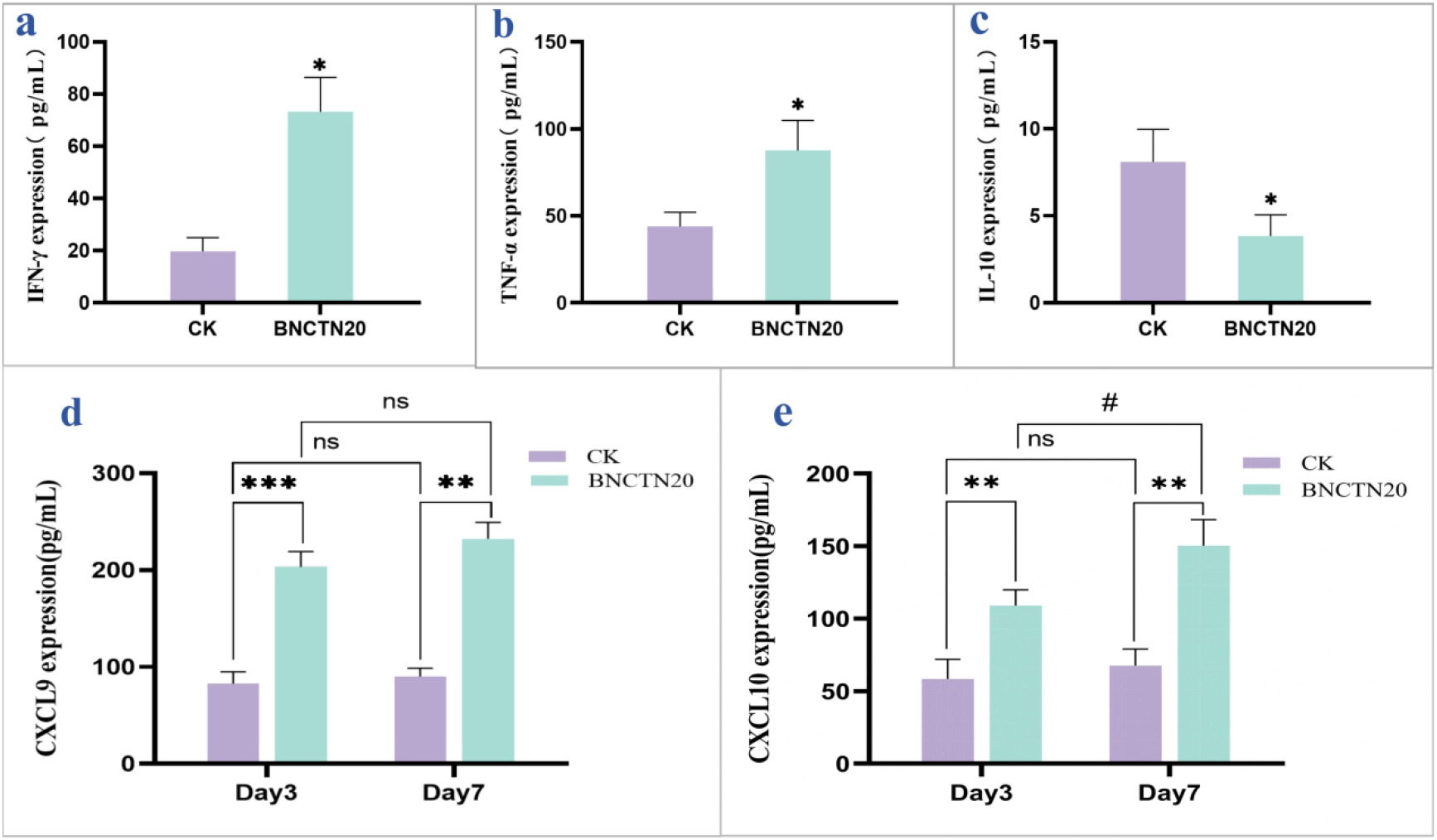
BNCT induces a systemic Th1 immune response and enhances CXCL9/CXCL10 chemotactic signaling. Two-group comparisons were performed using Welch’s t-test; multi-group comparisons were performed using one-way ANOVA with Tukey’s multiple comparisons test. (a) IFN-γ. (b) TNF-α. (c) IL-10. (d) CXCL9. (e) CXCL10.

Further analysis of the dynamic expression of CXCL9/10 on days 3 and 7 revealed distinct temporal kinetics. CXCL9 reached a high level as early as day 3 after irradiation and was maintained through day 7 (2.46-fold higher than CK, *p* < 0.001), displaying a rapid-response–plateau pattern (Fig. 4d). In contrast, CXCL10 increased progressively: 1.87-fold above CK on day 3 (*p* < 0.01), expanding further to 2.22-fold on day 7 (*p* < 0.01), and within the BNCTN20 group, day 7 was significantly higher than day 3 (*p* < 0.05), displaying a late sustained-amplification pattern (Fig. 4e).

These results indicate that BNCT induces a Th1-biased systemic immune response and establishes temporally complementary CXCL9/CXCL10 chemotactic signaling through the early response of CXCL9 and the sustained elevation of CXCL10.

### BNCT Upregulates PD-L1 but Does Not Promote Intratumoral CD8⁺ T-Cell Infiltration

Quantitative immunohistochemical analysis on day 7 after irradiation showed that PD-L1 expression in tumors of the BNCTN20 group was significantly higher than in the control group, with a relative expression intensity 1.62-fold that of the control (*p* < 0.01; Fig. 5a).

**FIG. 5.**
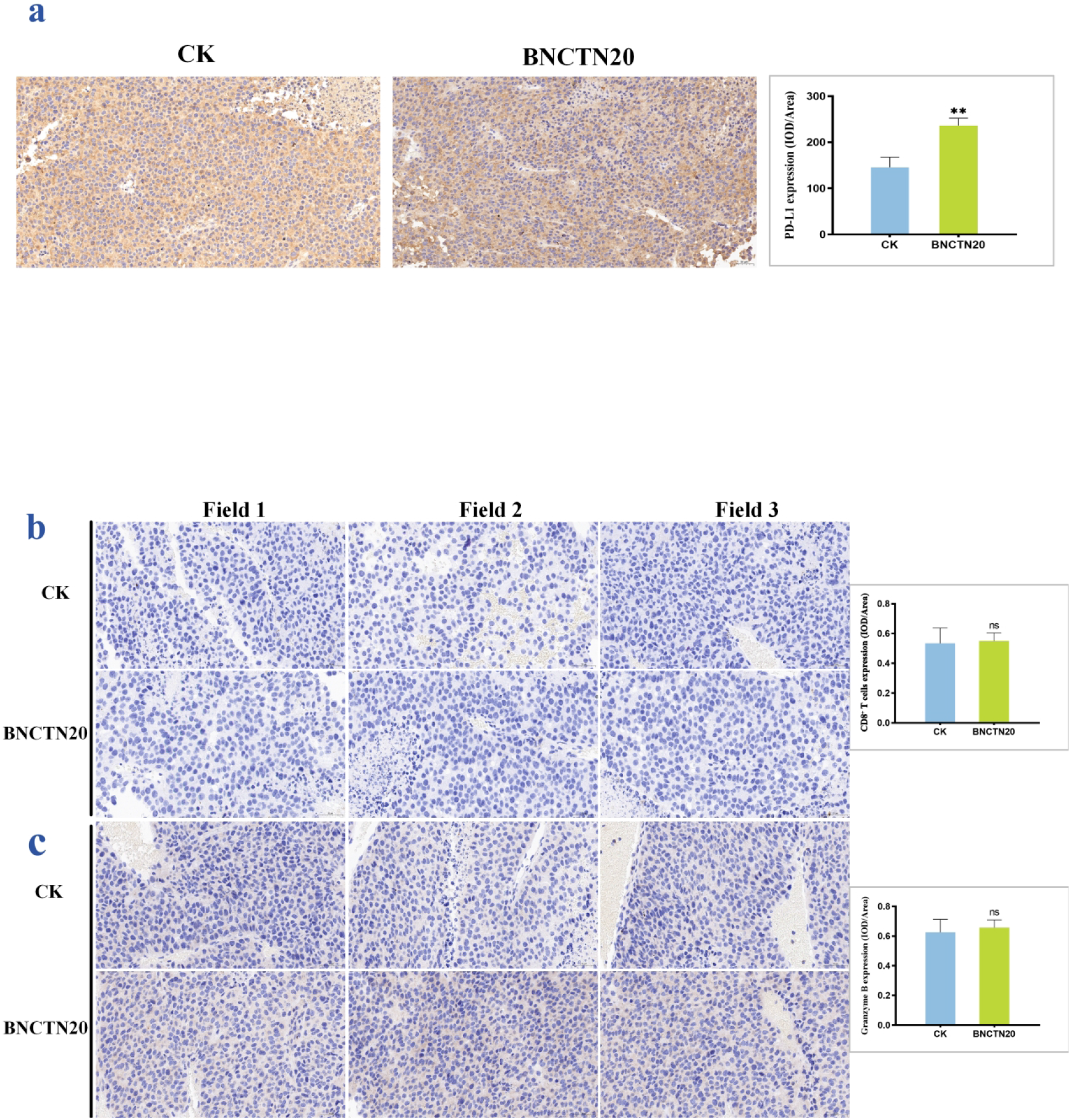
Effects of BNCT irradiation on PD-L1 expression and CD8⁺ T-cell infiltration in tumor tissue. (a) PD-L1 immunohistochemical staining on day 7 (brown-yellow: PD-L1-positive signal; blue: hematoxylin-counterstained nuclei) and quantitative analysis (p < 0.01). (b) CD8⁺ T-cell immunohistochemical staining images on day 7 (brown-yellow: CD8⁺ T cells; blue: hematoxylin-counterstained nuclei; multiple fields were observed per group, with no statistically significant intergroup differences; three representative fields are shown) and quantitative analysis. (c) Granzyme B immunohistochemical staining images on day 7 and quantitative analysis; multiple fields were observed per group, with no statistically significant intergroup differences; three representative fields are shown. Intergroup comparisons were performed using Welch’s t-test.

Notably, however, in contrast to the multiple positive findings of systemic immune activation described above, neither intratumoral CD8⁺ T-cell infiltration density nor granzyme B-positive expression increased significantly in the BNCTN20 group; semi-quantitative immunohistochemical analysis showed no statistically significant differences between groups (both *p* > 0.05; Fig. 5b, c).

### Boron Accumulation and Tissue Distribution of BPA in Bilateral Tumors

To evaluate the targeting selectivity of BPA in multiple tumor foci, ¹⁰B concentrations in blood, bilateral tumors, and major normal tissues at different time points after administration were measured by inductively coupled plasma optical emission spectrometry (ICP-OES). Concentration–time curves showed (Fig. 6a) that ¹⁰B concentrations in bilateral tumors peaked at 0.5–1 h after administration: at 1 h, the right dorsal tumor contained 29.28 ± 1.87 μg/g and the left shoulder tumor 26.86 ± 3.41 μg/g, both far exceeding the 20 ppm therapeutic threshold, with no significant difference between the two sides (*p* > 0.05). The concurrent blood concentration was only 7.85 ± 0.80 μg/g and was rapidly cleared over time. The tumor-to-blood (T/B) ratio peaked at 1 h after administration (right: 3.76 ± 0.52; left: 3.42 ± 0.33; >3 in all individuals) and then gradually declined, remaining above 2 at 7 h (Fig. 6b). Tumor-to-normal tissue (T/N) ratios at 1 h after administration showed (Fig. 6c) that the tumor-to-brain ratio was the highest (5.54 ± 0.66); ratios of tumor to liver, heart, lung, skin, and spleen ranged from 2.69 to 3.59; and the tumor-to-muscle ratio was 2.32 ± 0.35, all greater than 2. Only the tumor-to-kidney ratio was below 1 (0.83 ± 0.09), and the renal ¹⁰B concentration (34.00 ± 1.17 μg/g) was the highest among all tissues. None of the ratios differed significantly between bilateral tumors (*p* > 0.05).

**FIG. 6.**
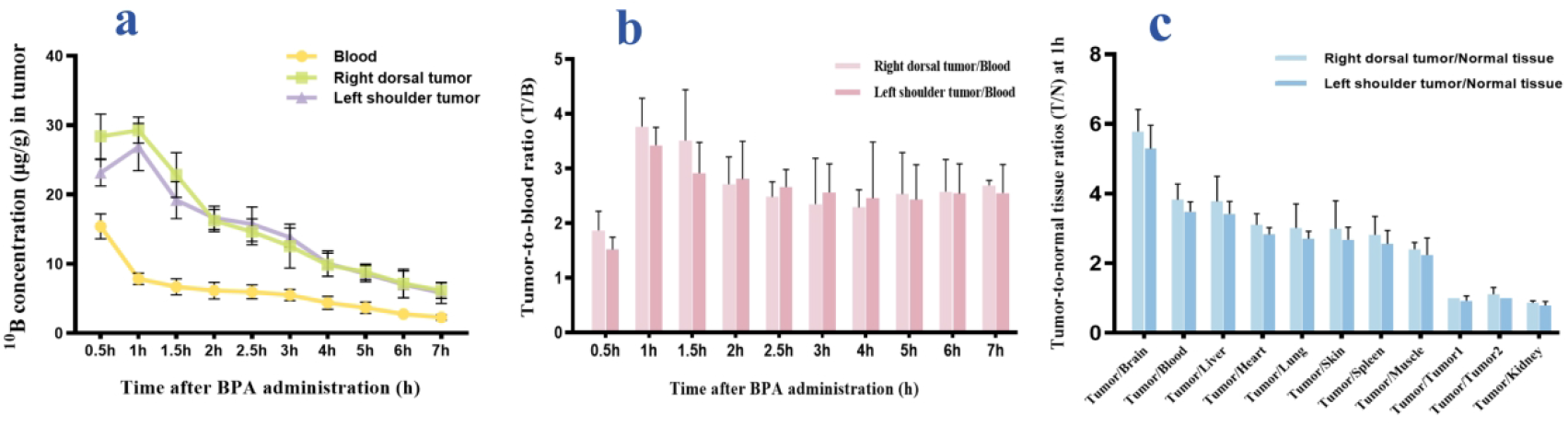
Biodistribution of boron in tumor-bearing mice after BPA administration (n = 4). (a) Concentration–time curves of ¹⁰B in blood, right dorsal tumor, and left shoulder tumor at different time points after administration (measured by ICP-OES; unit: μg/g, i.e., ppm). (b) Tumor-to-blood (T/B) ¹⁰B concentration ratios at different time points. (c) Tumor-to-normal tissue (T/N) ¹⁰B concentration ratios at 1 h after administration. Comparisons between bilateral tumors and their ratio indices were performed using paired t-tests (bilateral tumors and corresponding normal tissue samples were collected from the same mouse); no significant differences were observed between bilateral tumors.

## Discussion

In this study, we systematically evaluated the antitumor efficacy and immunological effects of BPA-based BNCT in the immunologically “cold” B16 melanoma model, yielding two principal findings. First, BNCT exerted potent direct killing: the BNCTN20 group achieved a tumor growth inhibition rate of approximately 88.9%, with median survival nearly doubled relative to controls (15 vs. 8 days). This killing was accompanied by well-defined immune-activating events—time-dependent accumulation of DNA double-strand breaks, caspase-dependent apoptosis, and nuclear-to-extracellular translocation of HMGB1—which together initiated immunogenic cell death (ICD), followed by a systemic Th1-type response characterized by elevated IFN-γ and TNF-α and reduced IL-10, as well as temporally ordered activation of the CXCL9/CXCL10–CXCR3 chemokine axis, indicating that BNCT can effectively initiate antitumor immunity even in poorly immunogenic tumors. Second, chemotactic signaling was established before local effector recruitment, while PD-L1 was synchronously upregulated, an intervenable node. Thus, BNCT drives a cold tumor into an initial activation state, providing a window for combination with checkpoint blockade.that is, BNCT can drive a “cold tumor” into an initial state of immune activation, providing a clear intervention window for subsequent combination with immune checkpoint blockade to convert the initiated ICD signal into substantive antitumor immunity and to expand local killing into systemic, durable antitumor immune responses.

### Direct Killing by BNCT and Initiation of ICD

Tumor volume in the BNCTN20 group was only 11.1% of that in the control group, markedly superior to neutron irradiation alone (N20; TGI ≈ 49.8%), confirming that its efficacy derives mainly from the high-LET particles generated by the ¹⁰B(n,α)⁷Li reaction rather than the neutron background dose^6,7^. The range of α particles and ⁷Li nuclei is only 5–9 μm, approximately the diameter of a single cell, which dictates that BNCT killing strictly depends on the selective enrichment of boron within tumor cells and confines cell death to the tumor locally. This is the physical basis for the subsequent DNA damage, apoptosis, and ICD events.

Sun et al. demonstrated in the B16F10 model that BNCT better preserves immune cells than X-rays and induces persistent, irreparable DNA double-strand breaks^9,13^. The γH2AX dynamics in the present study are consistent with this: the BNCT group was significantly higher than the control on day 3, the difference widened further by day 7, and within-group expression on day 7 exceeded that on day 3, showing time-dependent accumulation, whereas the control group showed no such change, suggesting that the clustered damage inflicted by high-LET particles exceeds the repair capacity of tumor cells. Meanwhile, cleaved caspase-3 rose to 2.64-fold that of the control, and H&E staining revealed the coexistence of necrosis and apoptotic cells, indicating spatial heterogeneity in the mode of cell death after BNCT: necrosis-like death predominates in regions of high dose deposition, whereas sublethally damaged cells enter the caspase-dependent apoptotic program^14^, and together these constitute the cytological basis for DAMP release^15^.

HMGB1 is one of the key danger signals of ICD: within the nucleus it participates in DNA damage repair, whereas once released extracellularly it acts as a DAMP that activates innate immune receptors^16^. In this study, HMGB1 signal increased to approximately 3.18-fold that of the control as early as day 3, with part of the signal located outside the nucleus, indicating nuclear-to-extracellular translocation and release. This is consistent with previous evidence that extracellular HMGB1 can serve as an early biomarker of BNCT efficacy^17^ and acts upstream of DAMP-driven immune activation in radiotherapy-induced ICD^18,19^. Thus, γH2AX → caspase-3 → HMGB1 constitutes a complete temporal sequence, suggesting that BNCT drives a mixed-mode immunogenic cell death through irreparable DNA damage, thereby completing the initiating phase of antitumor immunity.

### Systemic Th1 Response and Temporal Activation of the CXCL9/CXCL10 Chemokine Axis

The initiation of ICD was corroborated at the systemic level: after BNCT, serum IFN-γ increased 3.71-fold and TNF-α increased 2.00-fold, whereas the anti-inflammatory cytokine IL-10 decreased to 0.47-fold of the control, displaying a typical Th1-polarized pattern consistent with previous observations^9^.

Notably, CXCL9 and CXCL10 showed markedly different induction kinetics. Although both are IFN-γ-inducible CXCR3 ligands that should theoretically be driven by the same upstream signal, their behaviors clearly diverged: CXCL9 rose 2.46-fold by day 3 and entered a plateau (maintained on day 7, with no significant within-group difference), displaying a “rapid response–plateau maintenance” pattern; CXCL10, in contrast, continued to amplify (1.87-fold on day 3, further increasing to 2.22-fold on day 7, with within-group day 7 > day 3). This dissociation suggests that the two chemokines may originate from different cellular compartments: the rapid plateau of CXCL9 is more consistent with an immediate response of myeloid cells (e.g., cDC1 and macrophages) to IFN-γ, whereas the sustained amplification of CXCL10 may reflect a secondary response of stromal and tumor cells under persistent IFN-γ exposure^20,21^. Thus, the temporal relationship of “IFN-γ elevation first, followed by sequential chemokine activation” constitutes a complete signaling axis, consistent with clinical evidence that the chemokine microenvironment determines T-cell recruitment in melanoma^20,22^.

“Signal Established but Effector Absent”: A Multidimensional Explanation for the Lack of CD8⁺ T-Cell Infiltration

In stark contrast to the systemic activation described above, neither intratumoral CD8⁺ T-cell infiltration density nor granzyme B expression increased significantly. This negative result is the most central finding of the present study and can be interpreted from five dimensions.

First, the “presence” of chemotactic signals does not equate to “effective recruitment” of effector cells. Single-cell and spatial omics studies have made clear that CXCL9-driven promotion of T-cell infiltration depends on chemokine-producing cells (e.g., CCR7neg cDC1) forming effective cellular cluster niches with CD8⁺ T cells in space, rather than on diffuse chemokine elevation^21^; moreover, CXCL9⁺ macrophages must undergo a polarity switch with SPP1⁺ immune-exclusionary macrophages to build a T-cell-inflamed microenvironment^23^. What we measured here was serum chemokine levels, whose elevation does not guarantee the formation of a spatially ordered chemotactic gradient within the tumor; tumor-intrinsic T-cell exclusion programs, fibroblast–TGF-β physical barriers, abnormal tumor vasculature, and chemokine sequestration by the extracellular matrix can all produce a scenario in which “the signal exists but T cells cannot enter the tumor parenchyma”^24^.

Second, immune translation lags behind direct killing, and this temporal gap itself constitutes an intervention window. Post-irradiation changes in immune cell profiles and PD-L1 upregulation may be delayed until days 7–14^25^, whereas our mechanistic observations were made on day 7, coinciding exactly with the efficacy endpoint, that is, at the moment BNCT achieved maximal tumor debulking, immune translation had not yet been completed. Sun et al. observed increased CD8⁺ tumor-infiltrating lymphocytes at later time points in B16F10^9^, indicating that this translation is not absent but merely delayed.

Therefore, the mismatch of “fast killing, slow immunity” defines the value of combination therapy: intervening with immune checkpoint blockade within the window in which ICD signaling has peaked but infiltration has not yet been established could theoretically accelerate and amplify this delayed translation.

Third, inflammatory infiltration has occurred, but the infiltrating populations have not yet entered the cytotoxic phase. The sparse inflammatory cell infiltration seen on H&E may be dominated by non-CD8α⁺ populations such as macrophages, NK cells, or CD4⁺ subsets, thereby presenting a CD8-negative immunohistochemical phenotype. This composition of infiltrate suggests that BNCT-induced immune recruitment has already been initiated and that the immune composition of the tumor microenvironment has begun to loosen, with cytotoxic T cells potentially moving in gradually at later time points.

Fourth, immune activation in a cold tumor is more compelling. B16/B16F10 barely responds to PD-1 blockade alone^5,9^, making it a stringent model for screening immune-activating interventions. In this model, single-agent BNCT still induced a clear systemic Th1 response and chemokine axis activation, demonstrating that its immune-igniting capacity is sufficient to breach the first half of the low-immunogenicity barrier; the uncompleted second half—the conversion into T-cell infiltration—may reflect the raised time and signal thresholds imposed by this model (acting together with the second point above), and this is precisely where combination strategies come into play.

Fifth, type I interferon signaling provides a well-defined molecular target for next-step potentiation. Studies in the B16F10 model have shown that activating the cGAS–STING–type I interferon axis with Mn²⁺ significantly enhances BNCT-induced DC maturation and CD8⁺ T-cell responses^26^. This suggests that a druggable molecular switch may underlie the “ICD initiation → T-cell infiltration” bottleneck observed in this study, and that combining BNCT with immune adjuvants such as STING agonists has clear mechanistic rationale.

### PD-L1 Upregulation and the Formation of Adaptive Immune Resistance

Another key finding of this study is the upregulation of PD-L1 to 1.62-fold that of the control on day 7. The coexistence of an enhanced Th1 response and upregulation of an immunosuppressive molecule confirms the classic “adaptive immune resistance” mechanism: IFN-γ is the most potent inducer of PD-L1, upregulating tumor-cell PD-L1 via the JAK/STAT–IRF1 pathway^27^, such that immune attack itself paradoxically strengthens the tumor’s defenses^28,29^. In addition, irradiated tumor cells can directly upregulate PD-L1 via the ATR–Chk1–STAT3 pathway, suppressing antigen presentation and limiting immune priming^30^; the persistent accumulation of γH2AX indicates an activated DNA damage response, and the ATR–Chk1 pathway may likewise have contributed to PD-L1 upregulation in B16 tumors, consistent with the time course in which PD-L1 peaks at days 7–14 after radiotherapy^25^, a local immune barrier is erected in advance, before effector cells arrive.

The synchronous upregulation of PD-L1 happens to point to the target for breaking this bottleneck: although it is not the cause of absent infiltration, it foretells that late-arriving effector T cells may be neutralized as soon as they enter the tumor. Removing this barrier is precisely where the value of combining immune checkpoint blockade lies.

### Homogeneity of Boron Distribution and the Feasibility of Multifocal Treatment

The boron distribution results from the bilateral tumor model provide another layer of support for the above efficacy. At peak uptake, both tumors reached ¹⁰B concentrations far exceeding the therapeutic threshold, and all indices were highly homogeneous between the two sides. The tumor selectivity of BPA depends on LAT1 transporters highly expressed on tumor cells^31^; accordingly, it is reasonable to infer that bilateral syngeneic graft tumors have highly consistent LAT1 expression levels and functional states, which constitutes the molecular basis for the homogeneity of multifocal uptake. This finding has clear clinical relevance: melanoma frequently presents with multifocal and metastatic distribution, and if every lesion can obtain an equivalent boron load, a single BPA infusion combined with field-divided or whole-body intensity-modulated irradiation could theoretically achieve synchronous multifocal treatment without lesion-by-lesion dosing. Moreover, the tumor-to-blood ratio remained above 2 for several hours after administration, providing a favorable pharmacokinetic window for the longer operating time required for multifocal irradiation; the renal ¹⁰B concentration was the highest among all tissues, consistent with the known pharmacokinetic profile of renal BPA excretion, highlighting the necessity of renal dose monitoring in clinical application. At the same time, it should be noted that the tumor-to-muscle ratio was only about 2.3, and normal-tissue boron accumulation remains a factor limiting dose escalation; developing novel boron carriers with higher LAT1 selectivity is a direction for further widening the therapeutic window^32^.

### Clinical Translational Implications: Rationale for Combination with Immune Checkpoint Blockade

Previous studies have shown that the immune effects of single-agent BNCT are limited and can be fully unleashed only in combination with immune checkpoint blockade. Fujimoto et al. found in an advanced B16F10 dual-tumor model resistant to both radiotherapy and immunotherapy that BNCT alone suppressed only tumors within the irradiation field and anti-PD-1 alone was ineffective, whereas their combination (B-NIT) induced elevated serum HMGB1, increased intratumoral CD8⁺ T-cell infiltration, and abscopal effects, which were abolished by CD8 depletion^5^. Chiu et al. likewise demonstrated that BNCT combined with anti-PD-L1 further prolonged B16-F10 growth delay and enhanced intratumoral T-cell infiltration and activation compared with monotherapy^12^; Trivillin et al. similarly observed in a colon cancer model that regional and abscopal effects were produced only when BNCT was combined with immunomodulation^33^.

The above studies have demonstrated the value of combination therapy from different angles, but the details and limiting factors of immune activation after BNCT monotherapy remain to be explored. Taken together, the available evidence reveals an emerging common rule: the DAMPs and chemotactic signals released by BNCT must be freed from checkpoint suppression before they can be converted into T-cell infiltration and systemic antitumor immunity. Under monotherapy conditions, the present study observed the coexistence of “ICD initiation, Th1 activation, and chemokine axis establishment” with “absent local infiltration and PD-L1 upregulation,” providing mechanistic corroboration of this rule—BNCT accomplishes the “ignition” of the cancer-immunity cycle, while PD-1/PD-L1 blockade is responsible for “releasing the brake,” and the two act in synergy to drive the full operation of the immune response. Combined with the first phase I clinical trial of AB-BNCT for melanoma and other tumors, which achieved a 70% overall response rate^8^, clinical exploration of BNCT combined with ICIs is well justified.

## Study Limitations

This study focused on the initiation phase of ICD induced by single-agent BNCT and points the way for subsequent research; the following aspects await further work. First, the mechanistic cohort included few time points, as it was designed to capture the early dynamics of ICD; the late infiltration peak and PD-L1 changes were not covered. Second, CD8⁺ T cells were used as the core readout, and the lineage composition of the inflammatory cells seen on H&E (CD45, CD68, NK1.1, CD4/FoxP3, etc.) has not been typed. Third, the causal relationship between cytokine elevation and antitumor efficacy requires validation in functional experiments. Fourth, the subcutaneous B16 model differs from the clinical microenvironment, and extrapolation should be made with caution, although its low-immunogenicity background also makes the ICD signals captured here more discriminating. Fifth, serum chemokine levels are not fully equivalent to intratumoral chemotactic gradients, and spatial distribution evidence remains to be supplemented. Sixth, type I interferon and the cGAS–STING pathway were not directly examined, and the related inferences require experimental confirmation. All of the above have been incorporated into our follow-up plans, and the baseline data established in this study will inform time-point selection and efficacy interpretation in future combination experiments.

## Conclusions

This study demonstrates that BNCT exerts dual effects of potent killing and immune activation in the B16 cold tumor model: on the one hand, it achieves significant tumor control and survival benefit through selective killing by high-LET particles; on the other hand, it initiates ICD via DNA damage, mixed-mode cell death, and HMGB1 release, and induces a systemic Th1-type immune response and temporal activation of the CXCL9/CXCL10 chemokine axis. However, the establishment of chemotactic signaling failed to translate into local CD8⁺ T-cell infiltration, and the synchronous upregulation of PD-L1 indicates that adaptive immune resistance had formed in parallel. The high homogeneity of boron distribution in the bilateral tumor model further supports its multifocal application potential.

This study provides a direct mechanistic basis and preclinical foundation for strategies combining BNCT with PD-1/PD-L1 blockade.

